# Structural convergence for tubulin binding of CPAP and vinca domain microtubule inhibitors

**DOI:** 10.1101/2021.10.19.464980

**Authors:** Valérie Campanacci, Agathe Urvoas, Liza Ammar Khodja, Magali Aumont-Nicaise, Magali Noiray, Sylvie Lachkar, Patrick A. Curmi, Philippe Minard, Benoît Gigant

## Abstract

Microtubule dynamics is regulated by various cellular proteins and perturbed by small molecule compounds. To what extent the mechanism of the former resembles that of the latter is an open question. We report here structures of tubulin bound to the PN2-3 domain of CPAP, a protein controlling the length of the centrioles. We show that an α-helix of the PN2-3 N-terminal region binds and caps the longitudinal surface of the tubulin β subunit. Moreover, a PN2-3 N-terminal stretch lies in a β-tubulin site also targeted by fungal and bacterial peptide-like inhibitors of the vinca domain, sharing a very similar binding mode with these compounds. Therefore, our results identify several characteristic features of cellular partners that bind to this site and highlight a structural convergence of CPAP with small molecule inhibitors of microtubule assembly.

---

Microtubules are eukaryotic dynamic filaments made of αβ-tubulin heterodimers and involved in major cellular functions including ciliogenesis, intracellular transport and cell division. The dynamics of microtubules is regulated by several families of proteins (1). It is also perturbed by many small molecule compounds that target a growing number of tubulin binding sites (2, 3). Some of these compounds have proven to be useful to fight human diseases, in particular in oncology (4, 5).

Among these microtubule dynamics inhibitors, a diverse set of molecules including vinca alkaloids (6) and short peptides or depsipeptides (7) interact with the so-called tubulin vinca domain. This site is located at the interface between two tubulin molecules arranged in a curved protofilament-like manner (2, 8, 9). These compounds therefore interact with the longitudinal surface of both α- and β-tubulin. Doing so, they prevent protofilaments from adopting the straight conformation observed in the microtubule core, which accounts for their microtubule assembly inhibition. Whereas the α-tubulin subsite of vinblastine also binds to the N-terminal β-hairpin motif of stathmin proteins (9), whether other cellular tubulin regulators similarly bind to the vinca domain pocket of the β subunit is not known.

Centrosomal P4.1-associated protein (CPAP, also named CENPJ) is a 1338-residue long protein originally identified as a partner of the cytoskeletal protein 4.1R-135 (10). CPAP is however best characterized as a protein involved in centriole length regulation, allowing persistent slow growth of the centriolar microtubule plus end, CPAP overexpression leading to abnormally long microtubules (11–15). This activity can be recapitulated by a minimal construct comprising the tubulin-binding domain of CPAP, named PN2-3 and delineated as the CPAP 311–422 region (16), a microtubule-binding domain which is C-terminal to PN2-3 (17) and a dimerization motif (12). When isolated, PN2-3 was shown to prevent microtubule assembly through a ~76 residue-long region (16, 18). Using a tubulin-binding DARPin protein (19) as a crystallization helper, a partial structure of PN2-3 bound to tubulin was reported (12, 14). Because the N-terminal part of PN2-3 competes with the DARPin for tubulin binding, it was not visible in the crystal structure, and only about 15 residues of the PN2-3 C-terminal region, defined as the SAC domain (12), were traced bound to a β-tubulin site that corresponds to the outer surface of the microtubule.

Here, we report structures of tubulin bound to CPAP constructs as ternary complexes with α-tubulin specific αRep artificial proteins (20, 21). We show that part of the PN2-3 N-terminal region interacts with the β-tubulin longitudinal surface in a way very similar to that of vinca domain peptide ligands, therefore defining a type of cellular partners that binds to this site.

## Results

### Structures of tubulin-bound PN2-3 define the interaction of this protein with tubulin

We took advantage of the recently selected α-tubulin specific αRep proteins (20) to obtain several crystal forms of αRep–tubulin–CPAP ternary complexes. The CPAP constructs used were based on the 321–397 fragment (Fig. 1A). First, we determined a 2.7 Å resolution structure (Table S1; crystal form 1, with one complex per asymmetric unit) in which the CPAP residues 339 to 387 could be traced (Fig. 1B). These crystals were obtained with the iiH5 αRep. Surprisingly, in addition to the previously described iiH5–tubulin interaction (20), we identified a second tubulin-binding site for the αRep partner (Fig. 1B). In this structure, the CPAP 339-360 region folds as an α-helix which is connected by a linker, which does not interact with tubulin, to the C-terminal 371-387 SAC domain (Fig. 1B). Accordingly, the linker residues are among the least conserved ones of the CPAP PN2-3 domain from vertebrates (Fig. 1A). This linker is nevertheless reasonably defined in our structure, being stabilized by crystal contacts (see below).

**Figure 1.**
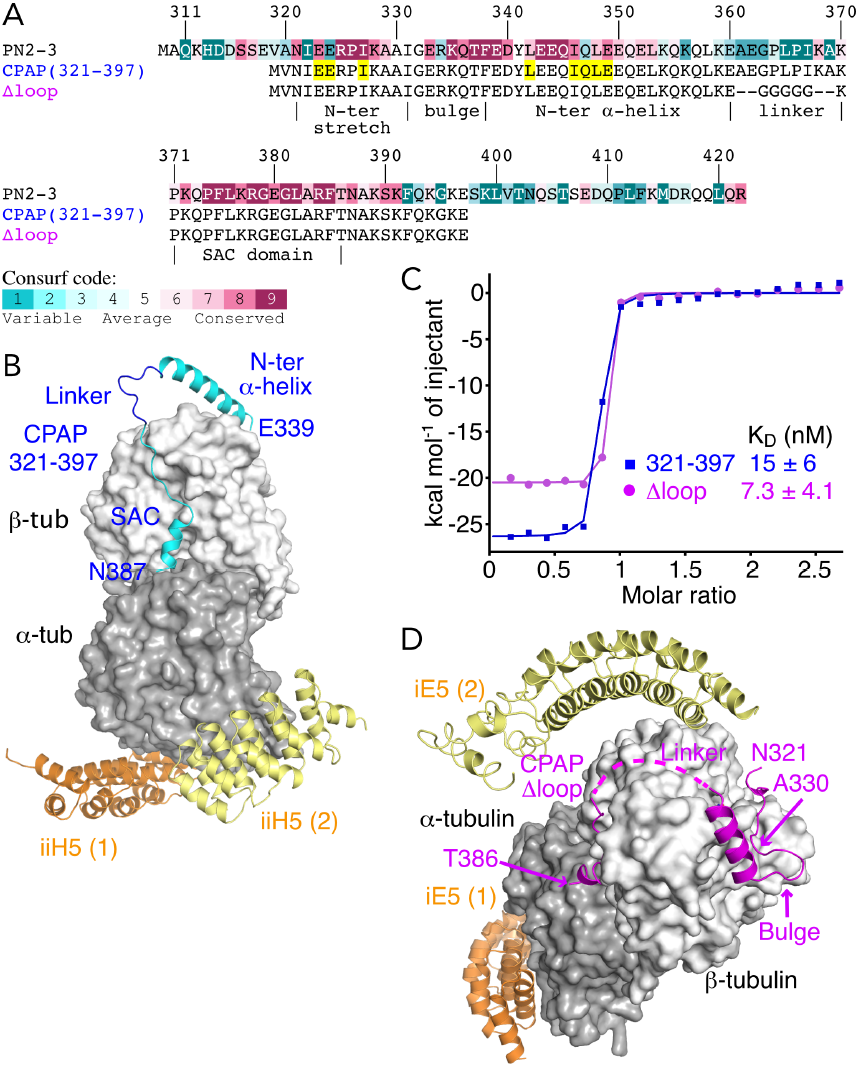
Overview of tubulin–CPAP structures. (**A**) Constructs used in this study and sequence conservation of the PN2-3 domain of CPAP. The PN2-3 sequence (residues 311–422 of CPAP, with an additional alanine after the N-terminal methionine) is aligned with that of constructs used in crystallization experiments. These constructs are based on the CPAP 321–397 region and include an additional valine after the N-terminal methionine. The residues 321 to 387 were traced in the different structures. This region has been divided into 5 structural motifs, as indicated under the sequence. Residues that have been mutated for affinity measurements are highlighted in yellow in the sequence of the 321–397 construct. The PN2-3 sequence conservation scores range from 1 (not conserved, blue) to 9 (highly conserved, magenta) according to the Consurf color code. (**B**) Crystal form 1: structure of the 321–397 CPAP fragment (cyan) in complex with tubulin (dark and light grey). Tubulin is further bound to two molecules (orange and yellow) of the iiH5 αRep used as a crystallization helper. (**C**) Isothermal titration calorimetry (ITC) of the interaction between tubulin and the 321–397 CPAP fragment or the Δloop construct (see also Fig. S1A,B and Table 1). (**D**) Crystal form 2: structure of tubulin bound to the CPAP Δloop fragment (magenta). The complex was crystallized with two molecules of the iE5 αRep. In panels B and D, the αReps in orange are those bound to tubulin in two structures of tubulin–αRep previously determined (20), the ones in yellow define new binding sites.

To enhance the likelihood to obtain other crystal forms, we replaced the linker region by a shorter 5-glycine motif (Fig. 1A). We checked by ITC the effect of this modification on the binding affinity for tubulin and found that the resulting “Δloop” CPAP construct displayed an affinity for tubulin similar to that of the unmodified protein (K_D_ about 7 nM and 15 nM, respectively; Fig. 1C, Fig. S1A,B, Table 1) and led to a high affinity complex. A new crystal form (Table S1; crystal form 2, two complexes per asymmetric unit), diffracting X-rays up to 2.35 Å resolution, was then obtained using the iE5 αRep crystallization chaperone. Remarkably, in this case too, two αRep molecules were bound per tubulin, identifying a second iE5 binding site in addition to the one previously described (20) (Fig. 1D), although the electron density was weak for the part of the αRep that did not interact with tubulin. In this structure, the 5-glycine linker was disordered. The PN2-3 N-terminal α-helix was also less well defined than in crystal form 1, its two C-terminal turns being disordered. In contrast, the region N-terminal to this helix became visible from residues Asn321 to Ala330, whereas residues 331 to 338 remained poorly defined. In the next sections, the information from these two structures will be combined for analysis. Information from two additional structures (see Methods and Table S1) will also be used to characterize the dynamic binding of the N-terminal region of the CPAP 321–397 fragment to tubulin.

**Table 1.** Thermodynamic binding parameters determined by ITC ^(a)^

| Construct <sup>(b)</sup> | K<sub>D</sub><br>(nM) | n | ΔH<br>(kcal mol<sup>-1</sup>) | -TΔS<br>(kcal mol<sup>-1</sup>) | ΔG<br>(kcal mol<sup>-1</sup>) |
| --- | --- | --- | --- | --- | --- |
| 321–397 (3) | 15 ± 6 | 0.79 ± 0.02 | -25.2 ± 1.0 | 14.6 ± 1.2 | -10.6 ± 0.3 |
| Δloop (9) | 7.3 ± 4.1 | 0.92 ± 0.22 | -19.1 ± 4.3 | 8.1 ± 4.1 | -11.0 ± 0.4 |
| LI>EE (4) | 65 ± 10 | 0.76 ± 0.04 | -22.8 ± 1.0 | 13.1 ± 1.1 | -9.6 ± 0.1 |
| QLE>GGG (5) | 47 ± 17 | 0.75 ± 0.10 | -20.7 ± 3.1 | 10.8 ± 2.9 | -10.0 ± 0.2 |
| I327R (3) | 449 ± 60 | 1.09 ± 0.14 | -17.5 ± 1.4 | 8.9 ± 1.4 | -8.5 ± 0.1 |
| I327W (4) | 29 ± 4 | 0.82 ± 0.01 | -21.7 ± 0.2 | 11.6 ± 0.3 | -10.1 ± 0.1 |
| EE>RR (4) | 34 ± 10 | 0.99 ± 0.22 | -19.0 ± 3.4 | 9.0 ± 3.3 | -10.0 ± 0.2 |
| 3R (4) | 275 ± 59 | 1.21 ± 0.04 | -14.2 ± 0.6 | 5.4 ± 0.6 | -8.8 ± 0.2 |

<sup>(a)</sup> mean ± s.d. <sup>(b)</sup> The number of measurements is indicated in parentheses.

### PN2-3 wraps around β-tubulin

Using αRep proteins as crystallization helpers, we were able to trace the entire region of PN2-3 predicted to interact with tubulin (18). This region comprises three modules, an N-terminal stretch, an α-helix and the SAC domain, that effectively interact with tubulin. They are separated by two non-interacting flexible modules, a “bulge” and the linker (Fig. 1, 2A,B). The bulge is poorly ordered, but the other modules are clearly defined in one structure or the other. PN2-3 wraps around the tubulin β subunit, with the stretch and α-helix N-terminal moiety that cap a β-tubulin surface which is engaged in longitudinal interactions within protofilaments. The flexibility conferred by the bulge allows the N-terminal stretch to be oriented about 90° to the α-helix and to point toward the lumen of the microtubule when PN2-3 is modeled at the plus end of a protofilament (Fig. 2A). Overall, these results provide a complete structural view of the binding of PN2-3 to tubulin, which agrees with previous in vitro competition and crosslinking experiments (12, 14, 18). They also provide a structural explanation for the inhibition of microtubule assembly by PN2-3.

**Figure 2.**
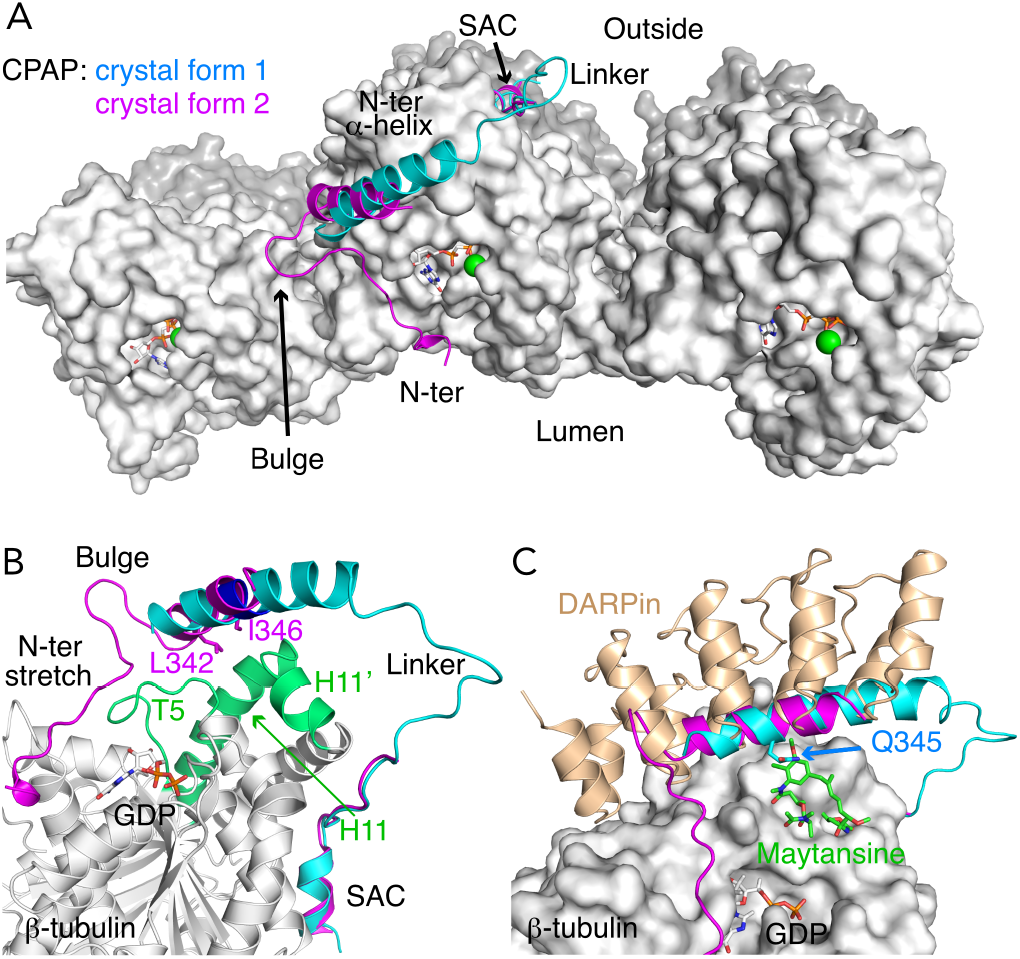
PN2-3 wraps around β-tubulin. (**A**) PN2-3 (cyan and magenta, from crystal forms 1 and 2, respectively) is modeled at the plus end of a microtubule (pdb id 6DPU), by superposing β-tubulin of tubulin–CPAP complexes to β-tubulin at the plus end of a protofilament. View from the plus end of the microtubule, three protofilaments are shown and CPAP is modeled on the middle one. The five structural motifs of PN2-3 are indicated (see also panel B and Fig. 1). (**B**) The PN2-3 N-terminal α-helix binding site. The β-tubulin elements interacting with this α-helix are highlighted in green. The side chains of residues Leu342 and Ile346, replaced in the LI>EE mutant, are shown in crystal form 2, whereas the main chain of residues 347 to 349, targeted in the QLE>GGG mutant, is in blue in crystal form 1. (**C**) Structural basis for PN2-3 competition with a DARPin (pdb id 5EIB) and with maytansine (4TV8) for tubulin binding. The side chain of Gln345, which would overlap with maytansine, is shown.

In its longest form, the PN2-3 N-terminal α-helix encompasses residues 339 to 360, in good agreement with the NMR-determined helical region of isolated PN2-3 (18). It binds in a groove at the tip of β-tubulin boxed in by loop T5 and the H11-H11’ region (Fig. 2B), giving a rationale for the competition for tubulin binding with the DARPin used as a crystallization helper in previous structural studies (12, 14) (Fig. 2C). It also explains the binding competition with maytansine (12, 22) (Fig. 2C). Only the N-terminal moiety of this CPAP α-helix interacts with tubulin. This part is the only one that is ordered in crystal form 2 (Fig. 2B). In addition to being two turns shorter, the α-helix in this crystal form is also rotated ~21° compared to crystal form 1 (Fig. 2A, Fig. S2A). These observations underline a dynamic binding of this α-helix to tubulin. This feature was further supported by the analysis of two other crystal structures of αRep–tubulin–CPAP complexes (Table S1), showing additional orientations of the helix (Fig. S2A). Moreover, inspection of the atomic temperature factors in the four structures indicated that those of the PN2-3 N-terminal region are higher than those of tubulin and of the CPAP SAC domain (Fig. S2B, Table S2). In addition, we studied the effect of two series of mutations in the N-terminal α-helix. We prepared the L342E-I346E (LI>EE) double mutant, targeting two hydrophobic residues pointing towards tubulin (Fig. 2B), and the ^347^QLE^349^>GGG triple mutant aiming to destabilize the α-helix. Both mutated proteins have an affinity for tubulin only slightly lower than that of the parental CPAP construct, as estimated by ITC (Table 1, Fig. S1C,D). Finally, we recorded the interaction of CPAP 321–397 with soblidotin-induced tubulin rings (8). Because PN2-3 inhibits the vinblastine-induced tubulin helical assembly (18), the CPAP construct was expected to inhibit also the formation of ring-type curved assemblies and it was indeed the case (Fig. S2C). Interestingly, when this construct was added to preformed rings, it did not induce their disassembly but interacted with them (Fig. S2C), although modeling indicated that the N-terminal α-helix of PN2-3 could not be accommodated at the tubulin longitudinal interface of curved assemblies (Fig. S2D). Taken together, these data argue for a moderate contribution of the PN2-3 N-terminal α-helix to the interaction with tubulin.

### The N-terminal stretch of PN2-3 targets the binding site of vinca domain inhibitors

The N-terminal stretch of the Δloop construct in crystal form 2 (residues 321 to 330 of CPAP) interacts with tubulin in an extended conformation, mostly without forming regular secondary structure apart a short 310 helix from residues 322 to 324. It binds in a site mainly contributed by helix H1, loop T5, and the H6-H7 region of the β subunit. It also interacts with the M-loop residue Arg278 and with GDP (Fig. 3A). Therefore, PN2-3 contacts the same β-tubulin structural elements as small molecule compounds binding to the vinca domain. Consistently, it has been shown that the PN2-3 affinity for tubulin drops ~14-fold in the presence of the vinca domain-targeting compound eribulin (12, 23). Strikingly, the PN2-3 N-terminal stretch closely overlaps with peptide and depsipeptide microtubule inhibitors, including phomopsin A, auristatins and tubulysins, four structural features being shared by CPAP and most of these compounds (24). They are described hereunder.

**Figure 3.**
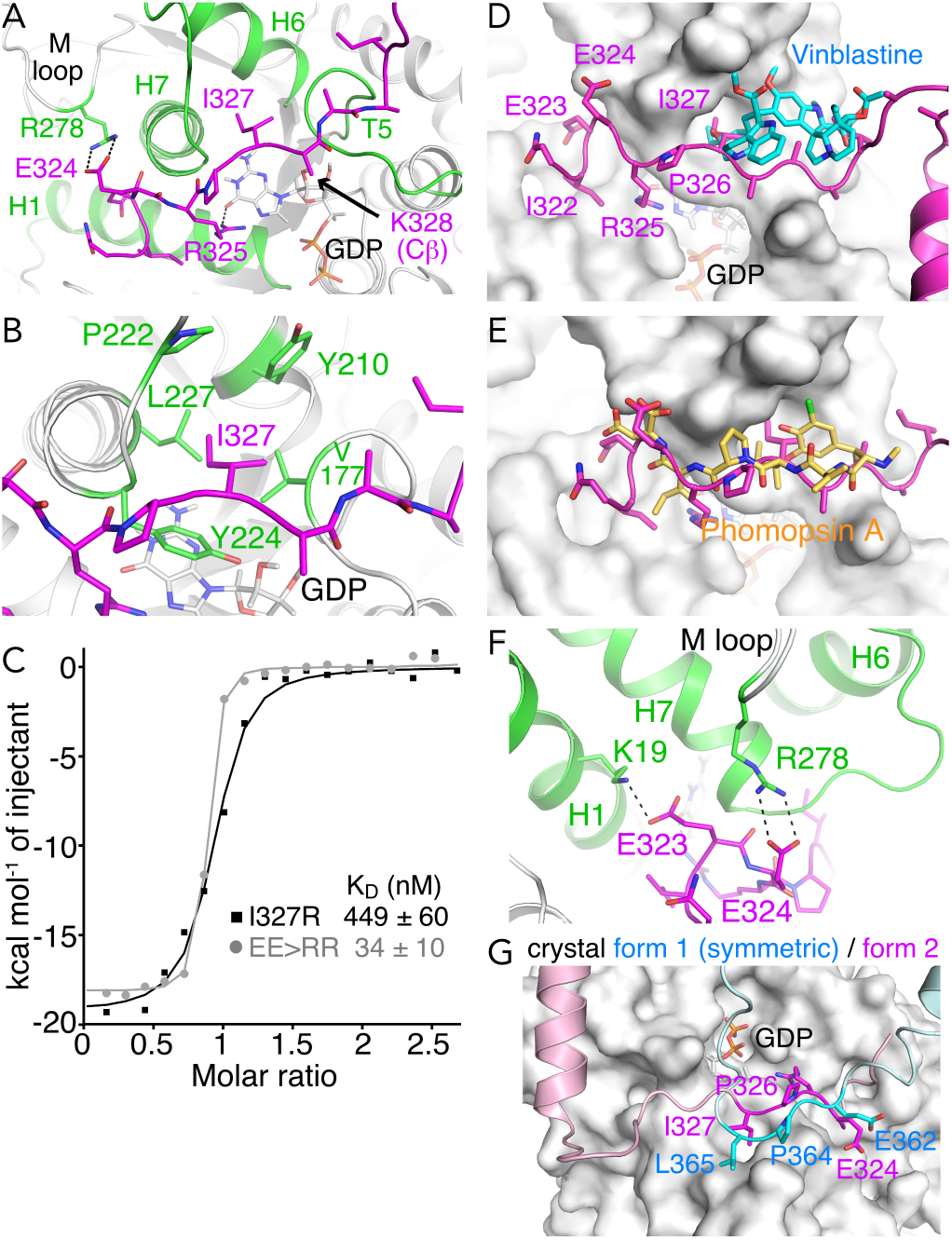
The interaction of the N-terminal stretch of PN2-3 with tubulin. (**A**) Overview of the interaction. The PN2-3 Δloop construct (from crystal form 2) is in magenta, and the tubulin-interacting elements are in green. The side chain of CPAP Lys328 which is weakly defined in the electron density maps has been truncated after its Cβ atom. (**B**) Zoom in on the interaction with tubulin of the CPAP residue Ile327. (**C**) ITC analysis of the tubulin:I327R and tubulin:E323R-E324R (EE>RR) interactions (see also Fig. S1E,G and Table 1). (**D,E**) The binding sites of vinblastine (pdb id 1Z2B, panel D) and of phomopsin A (3DU7, panel E) overlap with that of the N-terminal stretch of PN2-3. (**F**) Zoom in on the interaction with tubulin of the CPAP residues Glu323 and Glu324. (**G**) In crystal form 1, the linker region of a symmetry-related CPAP molecule (cyan) occupies the tubulin binding site of the N-terminal stretch of PN2-3.

First, the highly conserved Ile327 CPAP residue (Fig. 1A) points in a hydrophobic pocket of β-tubulin boxed in by Val177 (from T5), Tyr210 (H6), Pro222 (H6-H7 loop), Tyr224 and Leu227 (H7) (Fig. 3B). We tested the effect of an I327R substitution in the 321–397 construct and found a ~30-fold decrease of the binding affinity for tubulin (Fig. 3C, Fig. S1E, Table 1). This tubulin pocket is targeted by all the vinca domain ligands whose interaction with tubulin has been structurally characterized (Fig. 3D,E, Fig. S3). Remarkably, it accommodates a small aliphatic group in all cases, and it has been shown that replacing the corresponding isopropyl group in tubulysins by a bulkier aryl or cyclohexyl one leads to inactive compounds (25). However, in the case of CPAP, introducing the I327W mutation in the 321–397 construct did not decrease the affinity for tubulin (Table 1, Fig. S1F), indicating that, in the context of PN2-3 and in contrast to smaller peptide-like vinca domain inhibitors, a hydrophobic group bulkier than an Ile side chain is allowed at this position.

A second feature is the proline residue adjacent to Ile327 (Pro326, also conserved). A proline or proline-like residue is found at this position or at an adjacent one in most of the peptide-like inhibitors of tubulin (Fig. S3). Thirdly, whereas the Arg325 guanidino group interacts with the nucleotide base (Fig. 3A), its side chain aliphatic part together with Ile322 forms a hydrophobic patch interacting with β-tubulin, including with Tyr224. This tyrosine is in a stacking interaction with the guanine of the nucleotide and is involved in the regulation of microtubule assembly (26). Whereas the hydrophobic nature of the tubulin region interacting with these two CPAP residues is not strong, the observation that the vinca domain peptide inhibitors all have a hydrophobic group superposing with Ile322 and with the aliphatic part of Arg325 (Fig. 3E, Fig. S3) suggests such groups are important for tubulin binding. Accordingly, removing the corresponding phenyl group in tubulysin D leads to a compound ~40-fold less efficient in inhibiting cancer cell growth (27).

Finally, two acidic CPAP residues (Glu323 and Glu324) are in positions to make salt bridges with Lys19 and Arg278 of β-tubulin, respectively (Fig. 3F), suggesting that these glutamic acid residues contribute substantially to the interaction with tubulin. This proposal is supported by the observation that several small peptide-like compounds also have acidic groups overlapping with Glu323 and Glu324 (Fig. 3E, Fig. S3), and the interaction with the M-loop residue Arg278 has been proposed to enhance the affinity of some inhibitors for tubulin (28). In addition, several compounds shown to also bind to the “partial” binding site of the tubulin–stathmin complex, contributed by β-tubulin only, have one or two acidic groups (e.g. phomopsin A (8)). On the other hand, replacing the acidic group in tubulysins with, e.g., an hydrazine counterpart has only a mild effect on the activity (29, 30). In the case of CPAP, an E323R-E324R (EE>RR) mutant displayed an affinity for tubulin similar to that of the wild type construct (Fig. 3C, Fig. S1G, Table 1). Consistently, the affinity of a 3R mutant, having in addition the I327R substitution, was similar to that of the single I327R mutant (Fig. S1H, Table 1). It should also be noted that, whereas Glu323 is a conserved CPAP residue, Glu324 is not (Fig. 1A). We therefore conclude that Glu323 and Glu324 do not contribute much to the binding affinity of PN2-3 for tubulin.

Interestingly, whereas the N-terminal stretch of the 321–397 construct was not visible in crystal form 1, the vinca domain peptide site is not empty in that structure but occupied by the linker (which connects the N-terminal α-helix to the SAC domain) of a crystal-related symmetric molecule. Remarkably, the ^362^EGPL^365^ motif of the linker in that crystal form fills the space occupied by ^324^ERPI^327^ in crystal form 2 (Fig. 3G). This observation accounts for the destabilized N-terminal stretch in crystal form 1. More importantly, it confirms features important for targeting the vinca domain peptide of tubulin (Fig. S3).

## Discussion

In this study, we have shown that the PN2-3 domain of CPAP interacts with tubulin through three discrete, mostly linear binding motifs, a feature of intrinsically disordered proteins (31, 32). The overall affinity of PN2-3 for tubulin probably comes from the cumulative effect of these three interacting motifs, since constructs including either the N-terminal stretch and the N-terminal α-helix or the C-terminal SAC domain display K_D_ in the μM to the tens of μM range, i.e. two to three orders of magnitude higher than that of PN2-3 (12, 14). Such a binding mode, characterized by a cumulative contribution of weakly interacting points to the affinity, is likely to accommodate the mutation of (a priori) important residues. This proposal is supported by the moderate effect on the affinity of most mutations we studied, the most substantial being a 20- to 30-fold K_D_ increase with the I327R substitution. In contrast, the effect of similar modifications in shorter peptide-like inhibitors is more pronounced.

Another characteristic of intrinsically disordered proteins is their remaining conformational flexibility even when bound to their partners. Our results, which identify several orientations of the N-terminal α-helix (Fig. 2A, Fig. S2A) and an N-terminal stretch that is well defined in only one crystal form (Fig. 1B,D, Fig. S2B), indeed suggest that PN2-3 retains some mobility when bound to tubulin. This observation has implications for the mechanism of centriolar microtubule regulation by CPAP. Indeed, slow growth of the microtubule plus end that is observed in the presence of CPAP (12) implies that the N-terminal part of PN2-3 detaches from β-tubulin to unmask its longitudinal surface. The dynamic binding of this region to tubulin could lead to transient uncapping of protofilaments, therefore allowing microtubule elongation. Moreover, because CPAP constructs bind to tubulin rings (Fig. S2C,D), our results indicate that PN2-3 still interacts with tubulin even without the contribution of its N-terminal α-helix, consistent with the tracking by CPAP of the microtubule plus end during elongation (12, 14).

Another finding of our work which is relevant to the mechanism of centriole length regulation by CPAP is the orientation of the N-terminal end of PN2-3. When the tubulin–CPAP 321–397 construct structure is modeled at the plus tip of a protofilament, this end points to the lumen of the microtubule (Fig. 2A). Remarkably, it has recently been shown that the CP110–CEP97 complex binds to the luminal side of microtubule plus ends and that CP110 interacts with a CPAP region (residues 89 to 196) which is N-terminal to PN2-3 (33). Because CP110 also controls the length of the centrioles (11, 15, 33, 34), taken together, these data provide a basis for an interaction network at microtubule plus tips between centriolar microtubule regulators. To conclude, we have determined the complete structure of the microtubule destabilizing motif of the CPAP protein bound to tubulin. We have shown that its N-terminal stretch interacts with the vinca domain of β-tubulin. Its binding mode is very similar to that of several vinca domain peptide-like compounds from bacteria or fungi, which inhibit microtubule assembly. This finding points to a structural convergence for tubulin binding and provides constraints on the type of cellular partners accommodated in this pocket.

## Methods

### Proteins

The CPAP 321–397 construct was obtained by standard molecular biology methods from the PN2-3 DNA and inserted in the pET3d plasmid. Δloop and point mutants were then generated from this construct by PCR. All constructs were verified by sequencing. The corresponding proteins were overexpressed in the BL21(DE3) *E. coli* strain and purified as described for PN2-3 (18). Tubulin was purified from sheep brain (35). The iE5 and iiH5 αReps were produced and purified as described previously (20).

### Isothermal titration calorimetry (ITC)

Calorimetry experiments were conducted at 20°C with a MicroCal PEAQ-ITC instrument (Malvern). All proteins were buffer-exchanged to 20 mM Mes-K pH 6.8, 1 mM MgCl_2_, 0.01 mM EGTA and 0.01 mM GDP. Aliquots (2 μl) of CPAP constructs at 112 to 250 μM concentrations were injected into a 15 μM tubulin solution (cell volume, 0.24 ml). In the case of the Δloop construct, which has a higher affinity, titration experiments with 8 μM tubulin in the cell were also performed and led to a similar K_D_. However, because the parameter c=stoichiometry (n) • [tubulin]/KD remained slightly above the optimal conditions for an accurate estimation of the affinity parameters by ITC (36), the K_D_ value for this construct should be taken with caution. We also checked that the titration of buffer into 15 μM tubulin and that of the CPAP 321–397 and Δloop constructs into buffer yielded no heat signal (Fig. S1I-K).Analysis of the data was performed using the MicroCal PEAQ-ITC software provided by the manufacturer according to the one-binding-site model. The Origin software (Malvern) was used to superpose the data shown in Fig. 1C and 3C.

### Crystallization and structure determination

The αRep–tubulin–CPAP construct complexes were concentrated to about 20 mg ml^-1^ for crystallization experiments, leading to four crystal forms.

#### Crystal form 1

These crystals were obtained with the iiH5 αRep and with a CPAP 321–397 construct in which the lysine residues were methylated prior to the formation of the complex (37). In addition, residue Leu342 had been mutated to a methionine to help tracing the CPAP chain in lower resolution data using SeMet version of this construct at earlier stages of this project. Crystals were obtained at 293 K by vapor diffusion in a crystallization buffer consisting of 0.1 M Mes-K pH 6.8, 8 % (W/V) polyethylene glycol 3350. They were harvested in the crystallization buffer containing also 20% glycerol and flash-cooled in liquid nitrogen.

#### Crystal form 2

iE5–tubulin–CPAP Δloop construct crystals were obtained at 293 K in 0.18 M tri-ammonium citrate, 0.1 M Mes-K pH 6.8, 18.5 % (W/V) polyethylene glycol 3350 and cryoprotected in mother liquor supplemented with 20% glycerol.

#### Crystal form 3

The C-terminal tail of both tubulin subunits was cleaved by subtilisin (38) prior to the formation of a iiH5–tubulin–CPAP 321–397 construct complex. Crystals with one complex per asymmetric unit were obtained at 278 K in 0.2 M Na tartrate, 10% (W/V) polyethylene glycol 3350 and cryoprotected in mother liquor supplemented with 20% glycerol.

#### Crystal form 4

iE5–tubulin–CPAP 321–397 construct crystals (two complexes per asymmetric unit) were obtained at 293 K in 0.1 M Mes-K pH 6.0, 0.2 M Ca acetate, 20% (V/V) polyethylene glycol 400 and cryoprotected in the crystallization buffer containing 24% (V/V) polyethylene glycol 400.

Datasets were collected at 100 K at the SOLEIL Synchrotron (PROXIMA-1 and PROXIMA-2A beamlines). Data for crystal forms 1, 3 and 4 were processed with XDS (39) as implemented in the XDSME package (40). Crystal form 2 data were processed with autoPROC (41) which implements the STARANISO treatment for anisotropy (http://staraniso.globalphasing.org/). Structures were solved by molecular replacement with Phaser (42) using tubulin–iiH5 (pdb id 6GWD) and tubulin–iE5 (pdb id 6GWC) as search models, and refined with BUSTER (43) with iterative model building in Coot (44). In crystal forms 3 and 4, there was one αRep per complex, in contrast to crystal forms 1 and 2 which comprised an extra αRep (Fig. 1B,D). Data collection and refinement statistics are reported in Table S1. Figures of structural models were generated with PyMOL (www.pymol.org).

#### Sequence conservation

The sequence conservation of the CPAP PN2-3 domain (Fig. 1A) was calculated using the ConSurf program (45). The analysis was based on 150 sequences retrieved from the UniProt database and having a maximum of 95% sequence identity with human PN2-3 taken as a reference. About 40% of the sequences were from mammals, 55% from birds and 5% from reptiles, and there was one amphibian CPAP.

#### Data availability

Coordinates and structure factors have been deposited with the Protein Data Bank with accession numbers 7Q1E (crystal form 1), 7Q1F (crystal form 2), 7Z0F (crystal form 3), and 7Z0G (crystal form 4). Data supporting the findings of this study are available from the corresponding author upon reasonable request.

## Acknowledgements

We thank M. Knossow (I2BC, Gif-sur-Yvette) for many useful discussions and for a critical reading of the manuscript. Diffraction data were collected at SOLEIL synchrotron (PROXIMA-1 and PROXIMA-2A beamlines, Saint-Aubin, France). We are most grateful to the machine and beam line groups for making these experiments possible. This work has benefited from the crystallization and protein interaction platforms of I2BC supported by French Infrastructure for Integrated Structural Biology (FRISBI) ANR-10-INBS-05. Financial support by CNRS and by the Fondation ARC pour la Recherche sur le Cancer (Grants PJA20161204544 and ARCPJA2021050003651 to B.G.) is acknowledged.

