## Supplementary material for "Structural convergence for tubulin binding of CPAP and vinca domain microtubule inhibitors": Figures S1-S3, Tables S1-S2

### Supplementary Information

Figures S1 to S3, Tables S1 and S2, SI references

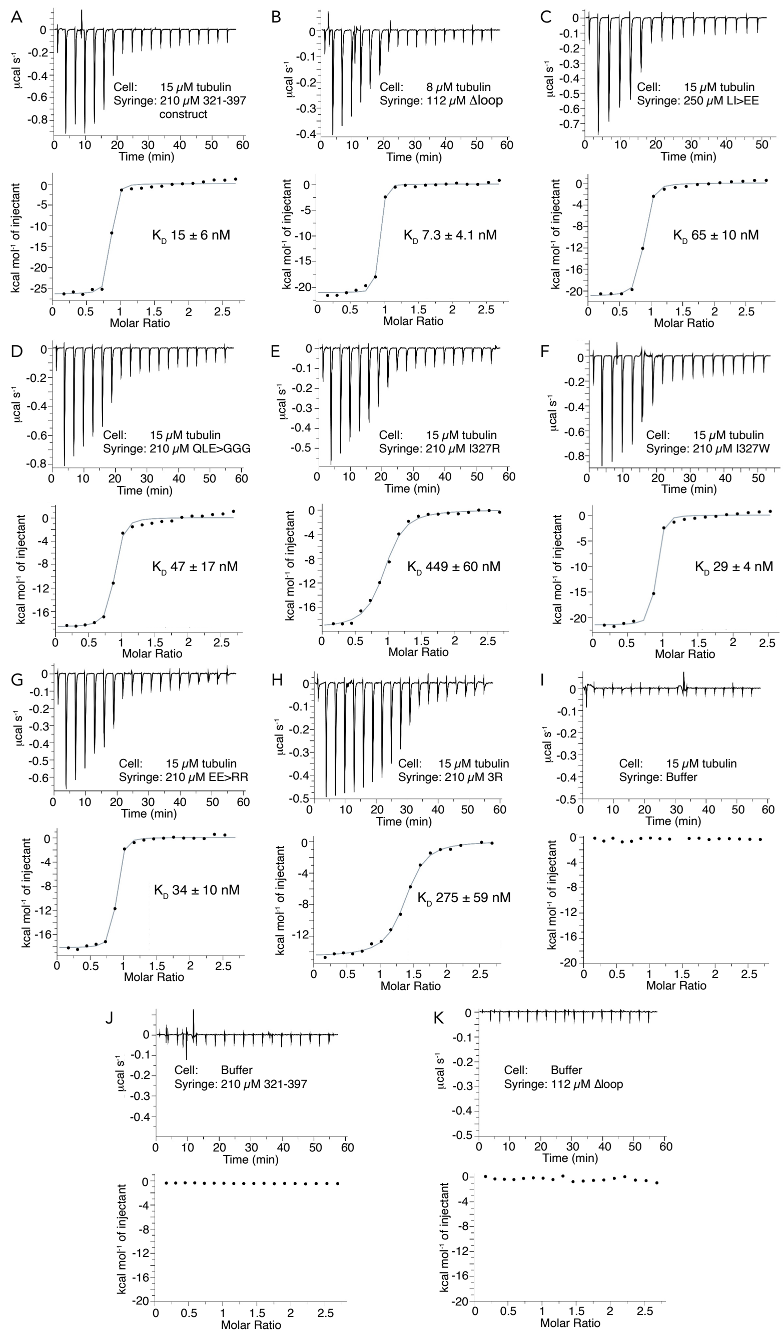


**Figure S1**. ITC analysis of the interaction between tubulin and CPAP constructs. (**A**) 321–397 construct. (**B**) ∆loop. (**C**) L342E-I346E "LI>EE" mutant. (**D**) $^{347}$QLE$^{349}$>GGG mutant. (**E**) I327R mutant. (**F**) I327W mutant. (**G**) E323R-E324R "EE>RR" mutant. (**H**) E323R-E324R-I327R "3R" mutant. Experiments were performed by stepwise titration of the CPAP construct into tubulin, at the indicated concentrations for the experiments shown. Upper panels display raw data; lower panels show the integrated heat changes and associated curve fits. The $K_D$ values were determined from at least three experiments (mean ± s.d.; see Table 1). (**I**) Titration of buffer into 15 μM tubulin. (**J**,**K**) Titration of the 321–397 construct (panel J) and of ∆loop (panel K) into buffer. The "Molar Ratio" in panels I to K is displayed as if there were 210 μM CPAP construct in the syringe (panel J) or 15 or 8 μM tubulin in the cell (panel J or K, respectively).

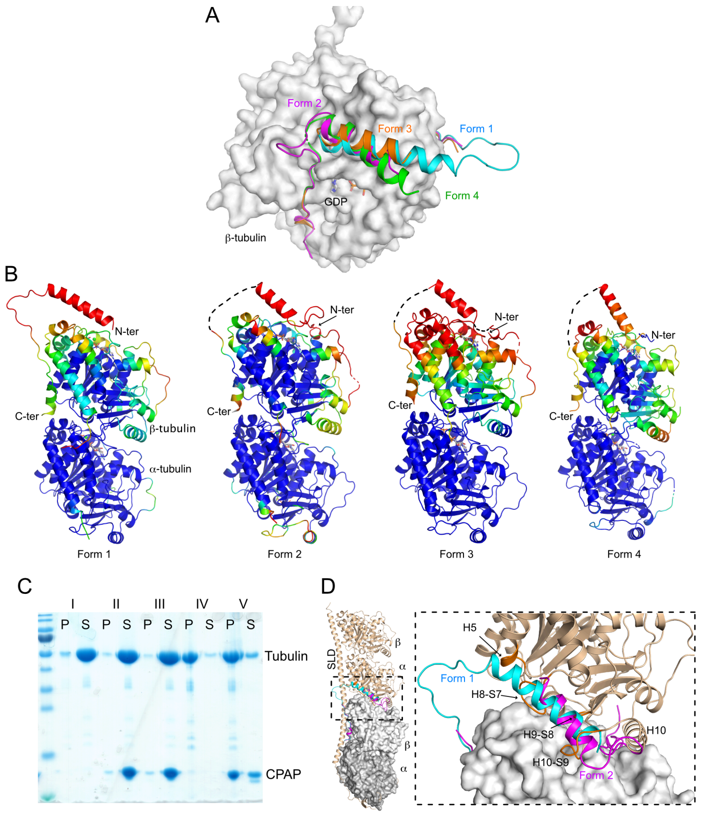


**Figure S2**. Analysis of the dynamic binding of the PN2-3 N-terminal region to tubulin. (**A**) Orientation of the N-terminal α-helix of PN2-3 in four αRep–tubulin–CPAP crystal structures. The tubulin β subunits have been superimposed. Taking the PN2-3 N-terminal α-helix of crystal form 1 (cyan) as a reference, that of the other crystal forms is misaligned by about 21° (form 2; magenta), 8.5° (form 3; orange), or 30° (form 4; green). (**B**) Temperature factors distribution in tubulin–CPAP structures (see also Table S2). The structures are rainbow colored according to temperature factors. Residues with a Cα temperature factor lower than the average temperature factor of tubulin–CPAP Cαs are in blue; residues with a Cα temperature factor higher than 1.5 times this average are in red. The N- and C-terminal ends of the CPAP fragment traced in the models are labeled; unmodeled regions within this fragment are shown as dashed lines. In crystal forms 3 and 4, electron density signal could be assigned to the N-terminal stretch of PN2-3 based on the form 2 structure, where this stretch is well defined. (**C**) Interaction of PN2-3 with tubulin ring aggregates. Tubulin (15 µM) was incubated at room temperature for 20 min in 20 mM Mes-K pH 6.8, 1 mM $MgCl_2$ in the following conditions: sample I, tubulin alone (control); sample II, tubulin + 30 µM CPAP 321–397 construct; sample III, same as sample II but 25 µM soblidotin (a dolastatin 10/auristatin analog) was added after 10 min incubation; sample IV, tubulin + 25 µM soblidotin; sample V, same as IV but 30 µM CPAP 321–397 construct was added after 10 min incubation. After high speed centrifugation (300,000 g for 15 min at 20°C), the supernatant (S) and pellet (P) (4 µg tubulin overall) were analyzed by SDS-PAGE on a Coomassie blue-stained gel. Whereas the CPAP construct prevents the soblidotin-induced tubulin aggregation (sample III), it hardly disassembles preformed ring aggregates but rather sediments with them (sample V). (**D**) Interference of the PN2-3 N-terminal α-helix with tubulin–tubulin longitudinal contacts in curved assemblies. The $T_2R$ structure, comprising two tubulin heterodimers and one stathmin-like domain (SLD) of the RB3 protein (pdb id 3RYC; Ref 1), is used as a curved tubulin assembly model. The β subunit of tubulin–CPAP (crystal forms 1 and 2) has been superimposed on that of $T_2R$ which interacts with the α subunit of the second tubulin of this complex. The N-terminal PN2-3 α-helix would clash in particular helix H5 and loops H8-S7, H9-S8 and H10-S9 of α-tubulin. (Left) Overview of the superimposed structures. (Right) Close-up of the part framed in the overview. For clarity, the α-tubulin C-terminal H12 helix and the SLD are not shown in the right panel. Tubulin–CPAP is colored as in Fig. 1, $T_2R$ is in wheat. Note that the tubulin–tubulin interface is expected to vary slightly between $T_2R$, which corresponds to a part of a 16-tubulin-containing ring or of an helical assembly having 16 tubulin molecules per turn (2, 3), and soblidotin-induced rings which contain 14 tubulin molecules (4). However, comparison with cryptophycin-induced 8-tubulin-containing rings (pdb id 7M18; Ref. 5) indicates that steric clashes would also take place in these smaller rings.

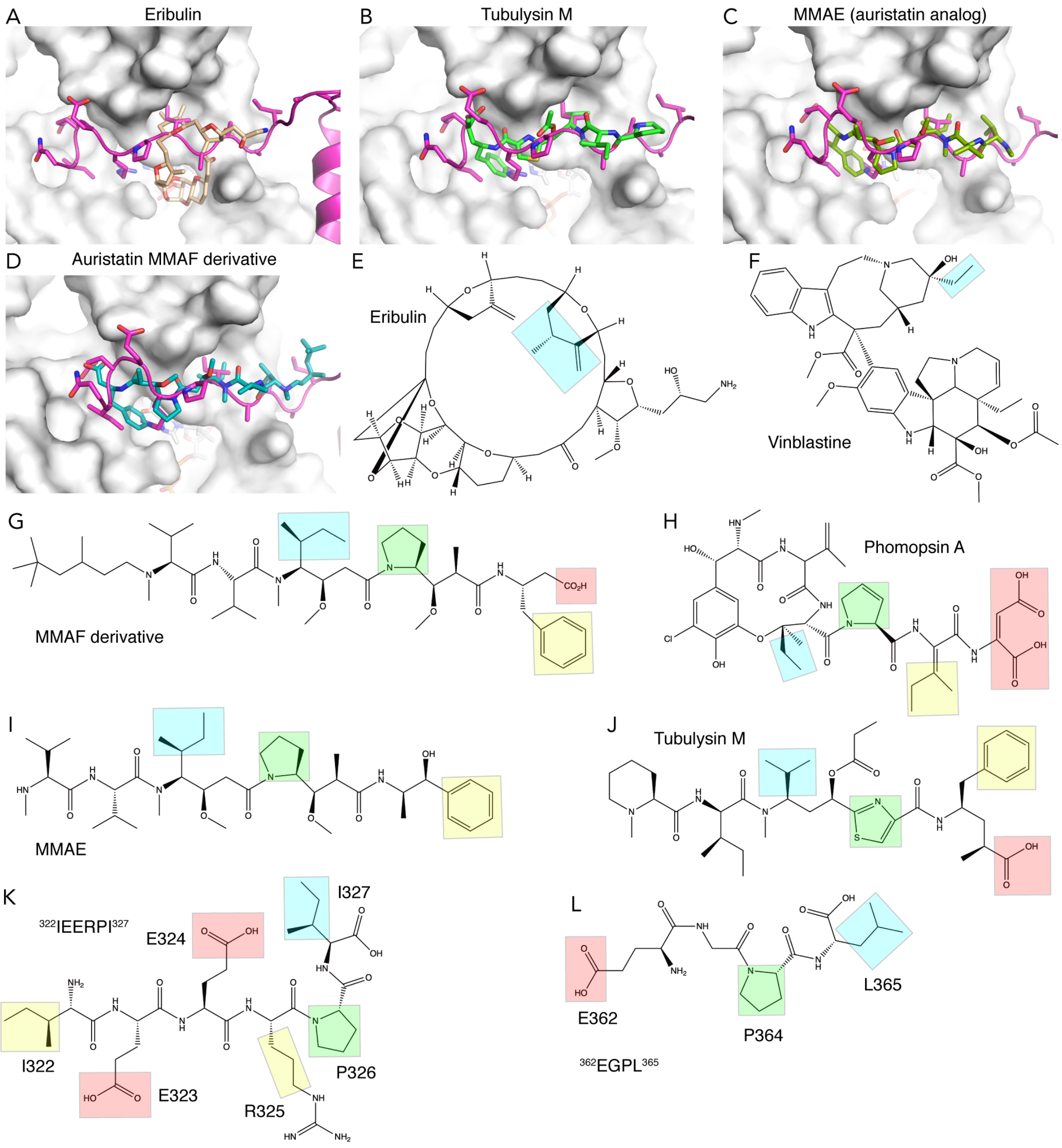


**Figure S3**. Comparison of the binding sites of vinca domain microtubule inhibitors with that of the N-terminal stretch of PN2-3. (**A**-**D**) Structures of tubulin-bound eribulin (pdb id 5JH7; Ref. 6), tubulysin M (4ZOL; Ref. 7), auristatin MMAE (5IYZ; Ref. 8), and an auristatin MMAF derivative (7JFR; Ref. 9) compared to that of the tubulin–Δloop complex (crystal form 2; the CPAP Δloop fragment is in magenta). (**E**-**L**) Chemical formulas of the compounds shown in panels A-D and in Fig. 3D,E, and of $^{322}$IEERPI$^{327}$ and $^{362}$EGPL$^{365}$ of CPAP. Highlighted are the parts of these molecules which overlap with Ile327 (cyan), with Ile322 and the aliphatic part of the side chain of Arg325 (yellow), and with the Glu323 and Glu324 acidic residues (red). Pro326 and Pro364 (panels K, L) and the proline-like groups in panels G–J are highlighted in green.

**Table S1. Data collection and refinement statistics.**

| | Form 1 | Form 2 | Form 3 | Form 4 |
|---|---|---|---|---|
| **Data collection**[a] | | | | |
| Space group | $P2_12_12_1$ | $P2_1$ | $P2_1$ | P1 |
| Cell dimensions | | | | |
| a, b, c (Å) | 53.6, 68.0, 420.9 | 89.1, 215.9, 95.5 | 54.3, 86.7, 137.6 | 57.1, 75.9, 151.3 |
| $\alpha, \beta, \gamma$ (°) | 90, 90, 90 | 90, 109.6, 90 | 90, 96.6, 90 | 96.2, 98.3, 111.6 |
| Resolution (Å) | 47.7-2.7 (2.8-2.7) | 108-2.35 (2.61-2.35) | 50-2.4 (2.48-2.4) | 50-3.49 (3.62-3.49) |
| Anisotropy resolution limits[b] | N.D. | 2.57, 2.84, 2.35 | N.D. | N.D. |
| $R_{meas}$ | 0.212 (5.39) | 0.189 (1.107) | 0.083 (1.556) | 0.272 (1.107) |
| $I / \sigma I$ | 12.23 (0.46) | 7.1 (1.6) | 13.06 (1.04) | 4.96 (0.97) |
| $CC_{1/2}$ | 0.999 (0.339) | 0.994 (0.675) | 0.998 (0.620) | 0.982 (0.394) |
| Completeness | | | | |
| spherical (%) | 99.9 (100.0) | 68.8 (12.7) | 99.2 (95.8) | 96.9 (84.4) |
| ellipsoidal (%) | N.D. | 90.8 (53.3) | N.D. | N.D. |
| Multiplicity | 35.6 (36.1) | 7.2 (6.8) | 4.5 (4.5) | 3.4 (2.0) |
| **Refinement** | | | | |
| Resolution (Å) | 2.70 | 2.35 | 2.40 | 3.49 |
| No. reflections | 43707 | 97303 | 49614 | 28469 |
| Rwork / Rfree | 0.200/0.254 | 0.188/0.220 | 0.221/0.272 | 0.221/0.266 |
| No. complexes/a.u. | 1 | 2 | 1 | 2 |
| No. atoms | | | | |
| Protein | 9200 | 19364 | 7868 | 16327 |
| Ligands | 72 | 191 | 67 | 146 |
| Solvent | 28 | 760 | 160 | 0 |
| B factors | | | | |
| Protein | 125.1 | 66.3 | 100.4 | 107.5 |
| Ligands | 109.4 | 62.1 | 107.7 | 116.3 |
| Solvent | 87.9 | 52.7 | 66.1 | N.A. |
| Coordinate error (Å) | 0.41 | 0.31 | 0.37 | 0.46 |
| R.m.s.d. | | | | |
| Bond lengths (Å) | 0.009 | 0.008 | 0.008 | 0.009 |
| Bond angles (°) | 1.00 | 0.96 | 1.00 | 0.98 |
| Ramachandran (%) | | | | |
| Favored region | 94.51 | 97.73 | 96.91 | 97.66 |
| Allowed region | 5.08 | 2.07 | 3.09 | 2.20 |
| Outliers | 0.41 | 0.20 | 0 | 0.14 |
| PDB ID | 7Q1E | 7Q1F | 7Z0F | 7Z0G |

[a] Data were collected on a single crystal. Values in parentheses are for the highest-resolution shell. [b] Determined by STARNISO using a local mean $I / \sigma(I)$=1.2. N.D., not determined. N.A., not applicable.

**Table S2**. Mean temperature factor of the Cαs of tubulin–CPAP in different crystal structures.

| | Crystal form 1 | Form 2 [b] | Form 3 | Form 4 [b] |
|---|---|---|---|---|
| Tubulin+CPAP [a] | 119 $Å^2$ (922) | 52 $Å^2$ (912) | 106 $Å^2$ (902) | 110 $Å^2$ (890) |
| α-tubulin | 101 $Å^2$ (440) | 45 $Å^2$ (432) | 76.5 $Å^2$ (438) | 93 $Å^2$ (434) |
| β-tubulin | 130 $Å^2$ (433) | 54 $Å^2$ (430) | 132 $Å^2$ (428) | 124 $Å^2$ (416) |
| CPAP | 175 $Å^2$ (49) | 100 $Å^2$ (50) | 164 $Å^2$ (36) | 151 $Å^2$ (40) |
| N-ter α-helix | 189 $Å^2$ (22) | 125 $Å^2$ (15) | 184 $Å^2$ (15) | 168 $Å^2$ (17) |

[a] Values in parentheses are the number of Cα atoms. [b] Analysis performed on the complex of the asymmetric unit for which the CPAP fragment is best defined in the electron density maps.
